# A de novo *GABPA* Variant in a Patient with Multifocal Cutaneous Vascular Tumors of an Unclassified Entity

**DOI:** 10.64898/2026.09.21.753178

**Authors:** Annegret Holm, Pascal Brouillard, Nagi Mahammmadzade, Mehrnaz Mehrabipour, Anna Shimbulu, Dionyssios Benetatos, Nisha Limaye, Ahmad Alomari, Leonard I. Zon, Whitney Eng, Harry Kozakewich, Friedrich G. Kapp, Miikka Vikkula, John B. Mulliken, Joyce Bischoff

## Abstract

We investigated the genetic basis of a previously unclassified congenital vascular anomaly in a patient with multifocal cutaneous vascular tumors. Through genomic analysis, we identified a novel heterozygous de novo germline variant in GA-binding protein-alpha (*GABPA*), an ETS family transcription factor. Histopathologic evaluation demonstrated capillary-venous lesions lacking glucose transporter 1 expression, distinguishing this entity from common infantile hemangioma. Notably, the tumor vasculature exhibited a prominent alpha-SMA-positive perivascular cell layer, with nuclear localized GABPA detected in endothelial and perivascular cells.

The *GABPA c.509T>G* (p.L170R) variant is absent from large population databases, including gnomAD, DeCAF, and RGC-MCPS databases, indicating it is extremely rare. The affected leucine residue is highly conserved across species and located within the Pointed (PNT) domain, a critical region mediating GABPA protein-protein interactions. AlphaMissense predicts this substitution to be highly damaging. Structure-based analysis using an AlphaFold-predicted approach, combined with in silico mutagenesis and molecular interaction analyses, revealed that the L170R substitution introduces new electrostatic interactions while disrupting native hydrophobic contacts within the PNT domain. Functional assessment in zebrafish demonstrated that mosaic stromal expression of GABPA-L170R caused abnormal vascular architecture compared with GABPA-wild-type controls, establishing a direct link between this variant and disrupted vascular development. Collectively, the genetic rarity, evolutionary conservation, predicted structural perturbation, and disruptive effects on vascular development establish *GABPA*-L170R as likely disease-causing variant. These findings define a new molecular mechanism underlying congenital vascular tumor formation and expand the role for ETS-family transcriptional regulation in human vascular anomalies.

## INTRODUCTION

Vascular anomalies encompass a heterogeneous group of vascular tumors and malformations (Goldenberg *et al*, 2025). The genetic basis of many of these disorders has been elucidated. However, atypical and unclassified presentations with unknown etiologies persist.

Vascular tumors range from benign (e.g., infantile and congenital hemangiomas, pyogenic granuloma), to borderline (e.g., Kaposi sarcoma, kaposiform hemangioendothelioma), to malignant (e.g., angiosarcoma) (Goldenberg *et al*., 2025). Some of these vascular tumor entities have documented genetic drivers. For example, most epithelioid hemangioendotheliomas arise from a gene fusion between endothelial expressed *WWTR1* and *CAMTA1*, resulting in inappropriate endothelial expression of CAMTA1 (Tanas *et al*, 2011). Potentially causative variants in *GNA11*, *GNA14*, and *GNAQ* have been reported in benign vascular tumors, such as congenital hemangioma (Jansen *et al*, 2021). In contrast, no causative variants have been reported for infantile hemangioma (IH) (Holm *et al*, 2024), in keeping with the relative paucity of genetic insights in vascular tumors compared to vascular malformations (Revencu *et al*, 2025; Seront *et al*, 2024). Germline variants with pathogenic significance in GA-binding protein alpha (*GABPA)* have not been reported, nor has a role for GABPA in vascular pathologies, including vascular tumors.

GABPA is a member of the E26 transformation-specific (ETS) family of transcription factors (Rosmarin *et al*, 2004). Encoded by the *GABPA* gene located on human chromosome *21q22.3*, GABPA contains a conserved ETS DNA-binding domain that recognizes purine-rich GA repeat sequences. Its Pointed (PNT) domain and an OST domain mediate protein-protein interactions – including binding with co-activators such as CBP/p300 – thereby facilitating assembly with other transcriptional regulators (Batchelor *et al*, 1998). GABPA associates with GABPB to form an αβ heterodimer and, on paired ETS-binding sites, can assemble into an α₂β₂ heterotetramer (Chinenov *et al*, 2000).

GABPA is essential for embryonic development, as murine homozygous null embryos do not survive to implantation (Ristevski *et al*, 2004). GABPA functions as a master regulator of gene expression for pluripotency during mouse preimplantation development (Zhou *et al*, 2025). At the cellular level, GABPA contributes to the transcriptional regulation of genes involved mitochondrial biogenesis (Yang *et al*, 2014) and cell cycle progression (Yang *et al*, 2007). GABPA further promotes tumor growth in malignant tumor entities, such as glioblastoma, melanoma, and hepatocellular and bladder carcinoma, by binding to mutated promoter regions of *TERT* (telomerase reverse transcriptase), resulting in TERT activation (Bell *et al*, 2015).

New studies have uncovered intriguing roles for GABPA in vascular development and endothelial specification. RNA sequencing of post-natal retinal vasculature and bioinformatic analysis of genome-wide promoter activity identified GABPA as a top transcriptional regulator of retinal vascular development (Jeong *et al*, 2017). A more recent study demonstrated that GABPA collaborates with the ETS pioneer transcription factor ETV2 to drive endothelial differentiation in human induced pluripotent stem cell-derived mesodermal progenitors, where both factors co-occupy newly accessible chromatin regions to enhance ETV2 transcriptional activity and trigger endothelial gene expression(Chen *et al*, 2025).

Herein, we report a de novo variant in *GABPA,* absent from large population databases, in a patient with dozens of cutaneous and subcutaneous congenital vascular tumors that were refractory to therapies used at the time for IH. Our in silico structural analyses showed that the introduction of a positively charged arginine (R) side chain in place of the hydrophobic leucine (L) side chain alters local interactions with neighboring residues. The L170R substitution favors a more stable complex between GABPA and the CBP/p300 transcriptional co-activator. Mosaic expression of the *GABPA* variant under the stromal *cxcl12b* promoter in zebrafish resulted in significantly increased numbers of malformed blood vessels, suggesting that the L170R variant has damaging effects on embryonic vascular development in a non-cell autonomous manner. Together, our discovery of a de novo germline *GABPA* variant in a patient with multifocal, benign (sub-)cutaneous vascular tumors establishes a novel clinical entity. These findings may provide crucial insights into the molecular drivers underlying vascular tumorigenesis, the interplay of tumor endothelium and its microenvironment, and the broader transcriptional role of GABPA in vascular pathologies.

## RESULTS AND DISCUSSION

### Clinical Presentation

Herein, we present a female patient with unclassified congenital vascular tumors. She presented at birth with multiple lesions of varying sizes, protrusions, and discolorations over the trunk, the extremities, external genitalia, the ears, and the lower lip while sparing the remaining face. Some lesions displayed intermittent external or internal hemorrhage (**Figure 1A**, **B**).

**Figure 1.**
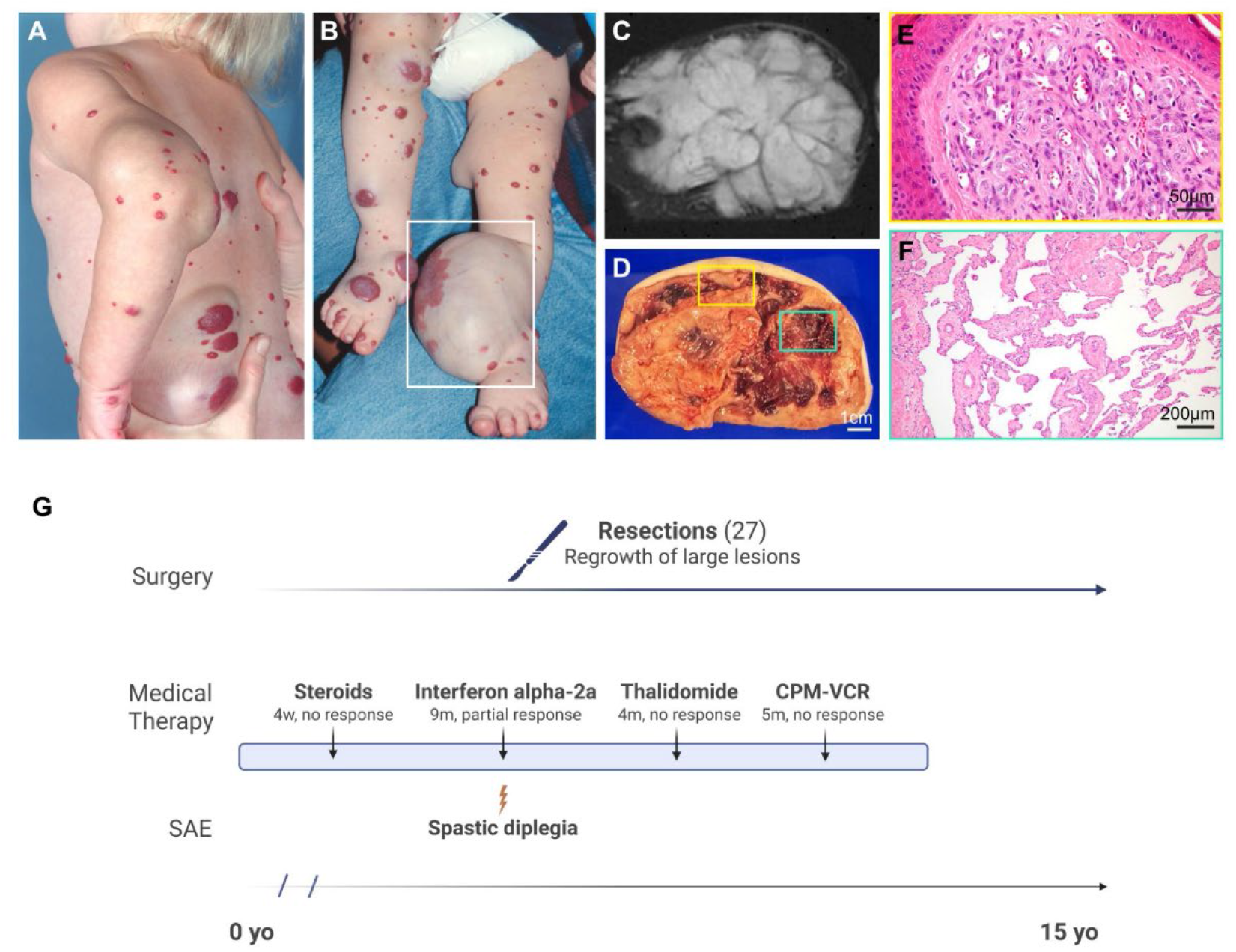
Clinical presentation, diagnostic workup, and therapeutic timeline of a congenital vascular tumor. (**A, B**) Clinical photographs showing diffuse, multifocal congenital cutaneous and subcutaneous vascular tumors extending across the upper extremities, trunk, and lower extremities. The white box in (B) highlights massive, localized lesion expansion over the left ankle. (**C**) Axial STIR MRI of the left lower extremity revealing a large, well-defined hyperintense soft tissue mass with prominent internal low-signal intensity septations. (**D**) Macroscopic cross-section of the resected ankle mass showing a heterogeneous tissue architecture with focal areas of hemorrhage and fibrous banding; colored insets frame regional variations chosen for microscopic evaluation. (**E**) High-magnification H&E-stained section corresponding to the superficial yellow box in (D), showing dense dermal lobules composed of small capillary-like channels lined by enlarged endothelial cells and surrounded by abundant pericytes (scale bar: 50 µm). (**F**) H&E-stained section corresponding to the green box in (D), exhibiting a distinct central zone characterized by large, anastomosing vascular channels bound by thin fibrous walls and flattened, inconspicuous endothelium (scale bar: 200 µm). (**G**) 15-year clinical treatment timeline illustrating a highly refractory course. Schematic charts the continuous requirement for multiple surgical interventions (27 total resections) with partial regrowth of large lesions. Arrows indicate sequentially failed medical therapies, including systemic corticosteroids, interferon alpha-2a (discontinued due to a severe adverse event [SAE] of spastic diplegia), thalidomide, and combination cyclophosphamide-vincristine (CPM-VCR) chemotherapy.

Initially assuming the patient’s vascular tumors may resemble IH biology, the patient started on corticosteroids one day postnatally, which was the mainstay of treatment for IH at that time. Because of persistent lesional growth while on corticosteroid therapy, the patient was switched to interferon-alpha-2a at 6 weeks of age (Ezekowitz *et al*, 1992). Increased doses over time resulted in stagnated growth and softening of most lesions. However, interferon-alpha-2a was discontinued after 9 months due to early signs of spastic diplegia. Further, regrowth and bleeding of larger lesions was noted towards the end of the interferon-alpha-2a treatment course. Sequential alternative medical treatment followed and included thalidomide, and a combination of cyclophosphamide and vincristine; again, with no significant clinical benefit for these medical treatments, while tolerated well (**Figure 1G**). In parallel and thereafter, the patient underwent multiple resections of large, protruding, functionally impairing, and in part bleeding lesions between ages 8 months to 15 years, comprising the following locations: upper and lower extremities including the elbow, flank, knee, and ankle region, the right posterior parietal to scalp region, the anterior and right neck, the auricular regions bilaterally, and the right flank (**Figure 1A**, **B**, **G**; **Supplemental Figure 1**). Endothelial cells were isolated from several resected lesions of the patient for in vitro studies (Boye *et al*, 2001) and DNA sequencing (**Supplemental Figure 2D**). Persistent regrowth of large lesions post resection required multiple operations. Most lesions eventually softened and underwent slow, spontaneous regression without medical treatment over the course of 16 years. While most lesions present at birth regressed, small and flat, bluish lesions appeared on the upper extremities during adolescence and adulthood; some of these punctate lesions remain into adulthood and are asymptomatic.

### Tumor Characterization

Radiological assessment by STIR MRI imaging was performed across multiple lesions. Axial imaging of the prominent lesion on the left ankle showed a large and well-defined hyperintense soft tissue mass with smooth, low-intensity internal septations dividing the mass into separate yet grouped entities by some thick septations, likely collagenous bands (**Figure 1C**).

The macroscopic cross-section of the tumor mass on the left ankle was heterogeneous with foci of hemorrhage and fibrous banding (**Figure 1D**). Histopathological evaluation with hematoxylin & eosin staining revealed vascular density and heterogeneity with a lobular pattern in most lesions. The lobules were composed of capillarous channels with a single layer of cuboidal endothelial cells with minimal pink cytoplasm, slightly enlarged nuclei, occasional mitoses, and one or two layers of mostly conspicuous pericytes (**Figure 1E**). Bigger lesions also exhibited this pattern but the lobules were larger, less distinctive as such, and evolved into anastomosing and thin-walled channels of greater caliber with flattened endothelial cells possessing smaller and often vertically oriented nuclei, sparse cytoplasm, and an often multilayered perivascular wall (pericytes and vascular smooth muscle cells), and a dense collagenous stroma, as depicted in the ankle lesion in **Figure 1D-F**. The tumor blood vessels expressed the canonical endothelial marker CD31 (**Supplemental Figure 2B**) and were negative for the lymphatic marker LYVE-1 (**Supplemental Figure 2C**, **D**). Glucose transporter-1 (GLUT1), a well-established marker used to distinguish IH from other vascular tumors and vascular malformations (North *et al*, 2000), was absent in tumor endothelium (**Figure 2A**, **B**). The lack of GLUT1 delineates the unclassified vascular tumor from common IH. A striking feature of the tumor vasculature was the multilayered perivascular cells visualized by anti-alpha-SMA staining, which was much less prominent in IH and normal human skin (**Figure 2C**, **E**, **G**). The average thicknesses of the alpha-SMA positive vessel layers were 8.8, 1.3, and 2.4 μm in the patient tumor, IH, and in skin controls, as quantified in **Figure 2D**, **F**, **H**. This suggests an alteration in the perivascular cell compartment in this unique vascular tumor. Clinical characteristics of patient tumors, IH, and normal skin controls from other patients are listed in **Supplemental Figure 1** and **Supplemental Figure 2A**.

**Figure 2.**
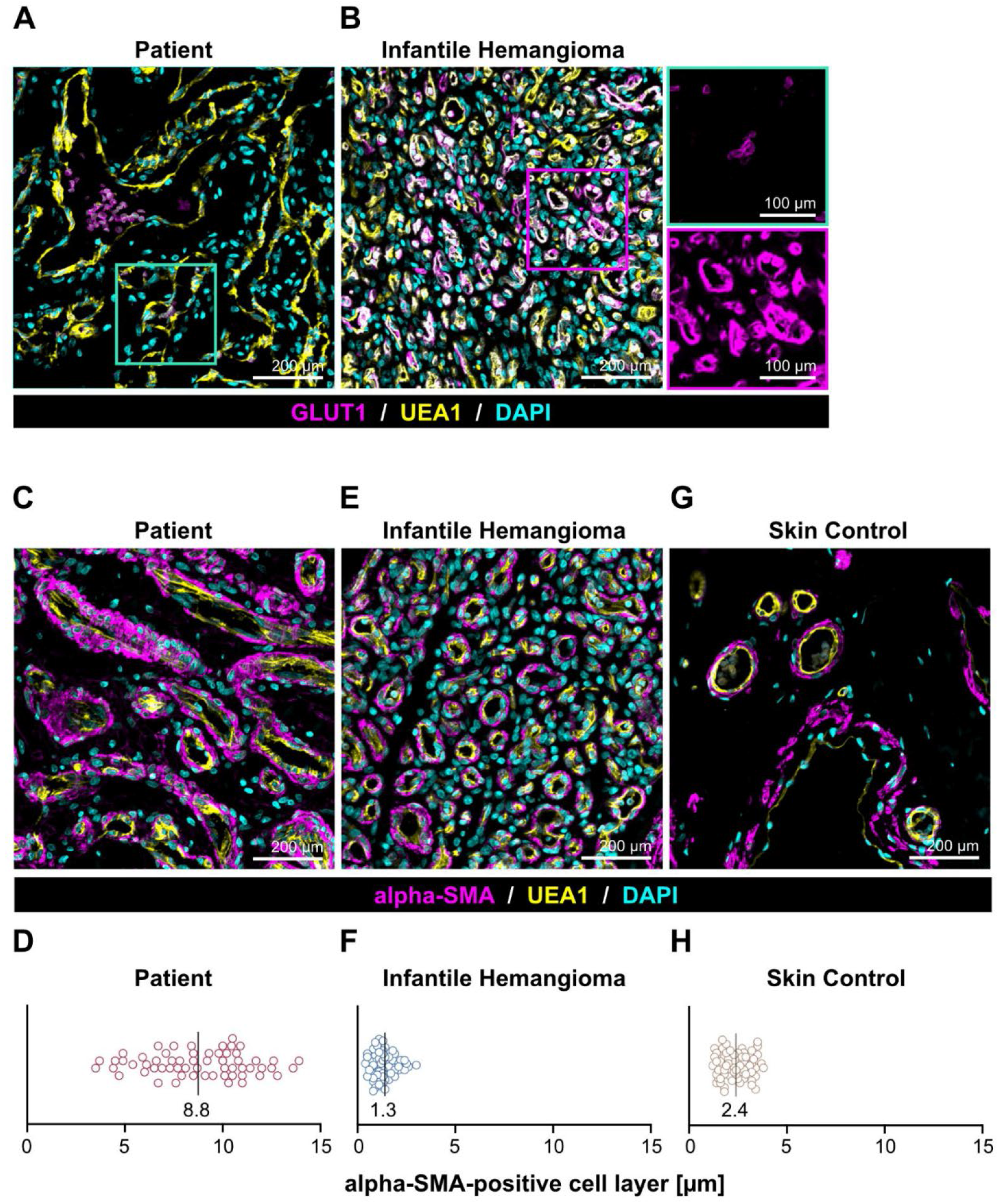
Immunohistochemical characterization of the tumor vasculature. (**A, B**) Immunofluorescence staining for glucose transporter 1 (GLUT1, magenta) counterstained with the endothelial marker *Ulex europaeus* agglutinin-1 (UEA1, yellow) and nuclear marker DAPI (cyan) in patient elbow (shown here), ankle and knee as well with positive controls from three individual IH patient samples. (**A**) GLUT1 was not detected in blood vessels in patient tumor tissue; higher magnification of the green boxed area (upper right panel) shows only faint, isolated signal within the vascular channels that represents natural autofluorescence of red blood cells. (**B**) Proliferating infantile hemangioma tumor tissue shows abundant GLUT1+ endothelium, serving as a positive control; the right bottom panel shows a higher magnification of the area highlighted by the magenta box (scale bars: 200 µm [main] and 100 µm [inset]). (**C**–**H**) Representative images of perivascular cell layer evaluated via alpha-smooth muscle actin (alpha-SMA, magenta) immunostaining paired with UEA1 (yellow) and DAPI (cyan). D, F, and H show corresponding measurement of perivascular layer thickness. (**C**, **D**) Patient’s lesional vessels (left elbow) have thickened layer of αSMA-positive perivascular cells, with a comparatively high average layer thickness of 8.8 µm. (**E, F**) Infantile hemangioma vessels show a compact perivascular wall with a lower average thickness of 1.3 µm. (**G, H**) Healthy, age-matched skin control tissue show perivascular layer thickness of 2.4 µm (scale bars for C, E, G: 200 µm; n=3 biological replicates for patient tissue [left elbow, knee, and ankle], infantile hemangioma, and skin control, respectively).

### Discovery of a de novo Variant in *GABPA*

Endothelial cells (EC) designated *HeomaEC-8* were isolated from vascular tumor tissue removed from different anatomical sites (Boye *et al*., 2001). Specific tumor locations of isolated HeomaECs are indicated by letters (**Supplemental Figure 3A**). Peripheral blood samples were obtained from the patient and her parents. Genomic DNA was isolated from patient tissue EC, denoted as Heoma-EC-8a, 8b, 8c, 8f, 8g, 8h, 8i, and peripheral blood mononuclear cells. A comprehensive next-generation sequencing panel (OncoPanel) did not identify genetic variants in the PI3K or RAS signaling pathways that are frequently detected in other vascular anomaly entities. Exome sequencing (ES) data from peripheral blood mononuclear cells from the trio was analyzed under dominant and recessive hypotheses. No strong candidate pathogenic variant could be identified. Using a de novo model, we discovered a likely pathogenic variant not inherited from the parents: *hg38_chr21_25752190T>G; c.509T>G*; L170R in the *GABPA* gene (**Figure 3A**). The variant is absent from population databases, including GnomAD_v4 (807,162 individuals), deCAF (150,119 UK genomes) and Regeneron Mexico City Prospective Study (MCPS; 150,996 individuals). It is predicted to be damaging by 11 out of 20 tools used in Highlander (http://sites.uclouvain.be/highlander/) and predicted to impact the protein structure by AlphaMissense (score 0.977; range 0-1) (Cheng *et al*, 2023). The patient appeared heterozygous (VAF range 35-55%) for the variant in peripheral blood, in lesional ECs (19), and tumor tissue from the elbow and knee (**Figure 3B**). Our study, however, does not differentiate between a true constitutional germline event and early somatic mosaicism. Molecularly, L170 is a highly conserved amino acid and structural analysis using Dynamut2 (Rodrigues *et al*, 2021) predicted ΔΔG^stability^ as destabilizing (-0.81 kcal/mol). A similar effect was predicted by mCSM-Stability (Pires *et al*, 2014) (-1.445 kcal/mol). In summary, L170 is highly conserved, the variant is exceedingly rare, and the substitution of arginine (R) for leucine (L) at position 170 is likely to affect GABPA conformation and function.

**Figure 3:**
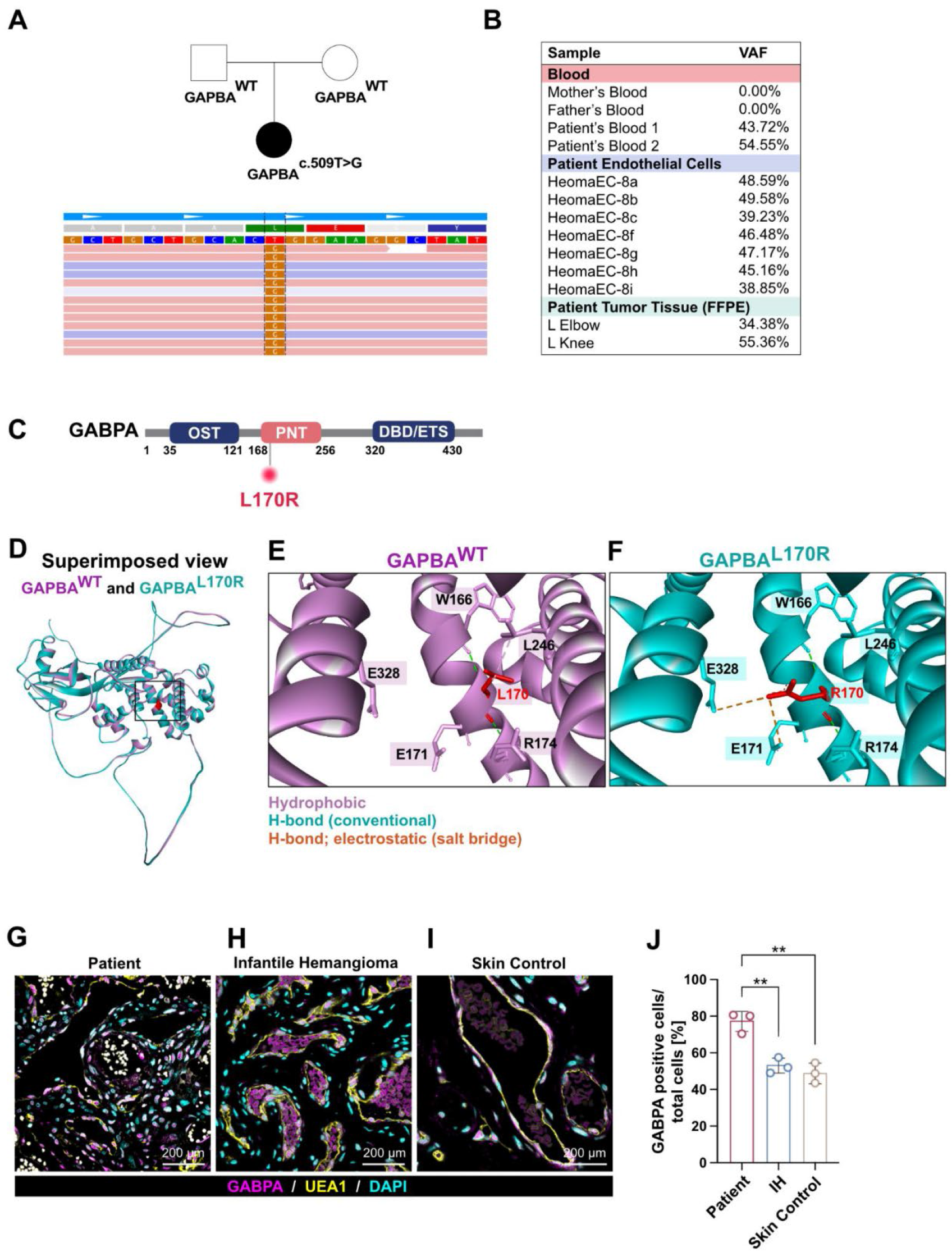
Identification of a *de novo* variant in *GABPA* (*c.509T>G*; L170R) in patient blood and tumor tissue. (**A**) Family pedigree showing the affected patient (black circle) carrying the *de novo GABPA c.509T>G* nucleotide substitution, and unaffected parents (white circle: mother; white square: father) are WT. Below, part of an alignment of forward (pink) and reverse (blue) reads showing T to G nucleotide change. (**B**) Table detailing variant allele frequencies (VAF). High VAF values (∼39% to 55%) are present across patient’s blood, isolated endothelial cells (*HeomaEC*) from different anatomical locations, and FFPE tumor tissue (**C**) Schematic of GABPA protein structural domains showing the location of the L170R variant. (**D**) Superimposed view of the AlphaFold-predicted GABPA wild-type structure (purple) and the in silico-generated GABPA-L170R mutant structure (cyan). (**E**, **F**) Zoomed in view of WT and variant. Hydrophobic interactions shown in purple; conventional H-bonds in green and electrostatic interactions in orange. (**G**-**I**) Immunofluorescent staining for wild-type GABPA (magenta), co-stained with UEA1 (yellow) and DAPI (cyan) in patient tissue (left elbow), infantile hemangioma, and age-matched skin control sections (scale bars: 200 µm). (**J**) Quantification of GAPBA positive cells shows significantly increased GAPBA in patient tumor tissue compared to infantile hemangioma and normal skin (p< 0.01). Notably, GABPA is expressed abundantly both in the endothelial and perivascular compartment in patient tissue. *n*=3 biological replicates: patient tissue from left elbow (shown), left knee and left ankle; IH and skin from 3 different individuals. One-way ANOVA with Šídák’s multiple comparisons test, data presented as mean ± SD.

### The *GABPA* L170R Variant Remodels Intramolecular Interactions

The L170R variant is located within the highly conserved PNT domain of GABPA (**Figure 3C**). Structural analysis revealed that substitution of the hydrophobic leucine residue with a positively charged arginine remodels the local intramolecular interaction network surrounding residue 170 without substantially altering the overall predicted protein fold (**Figure 3D**). In the wild-type structure, L170 participates in hydrophobic interaction with L246. In contrast, the L170R substitution abolishes this hydrophobic contact while preserving hydrogen-bond interactions involving W166 and R174. The introduced arginine establishes new electrostatic interactions with E171 within the PNT domain and E328 within the ETS DNA-binding domain (**Figure 3D-F**). These changes suggest that the L170R variant rewires intramolecular contacts and may alter communication between the PNT and ETS domains, potentially influencing GABPA conformational dynamics and downstream protein-protein interactions. A complete list of predicted intramolecular interactions identified for the wild-type and L170R structures is provided (**Supplemental data file 3**).

### Protein-Protein Interaction between GABPA PNT domain and CBP/p300

GABPA was first shown to bind directly to the transcriptional co-activator CREB binding protein (CREBBP or CBP/p300) in a study of IL-16 expression in T lymphocytes (Bannert *et al*, 1999). Subsequent studies demonstrated that the PNT domain in GABPA (amino acids 168-256) directly interacts with CBP/p300 in the context of retinoic acid induced myeloid cell differentiation (Resendes & Rosmarin, 2006). This interaction was further mapped to the CH3 domain (amino acids 1680-1850) of CBP/p300 and the PNT domain of GABPA (Kang *et al*, 2008).

To assess whether the L170R variant could influence this interaction, we performed protein-protein docking using full-length GABPA and a large CBP/p300 fragment encompassing residues 1323-1950, which includes the CH3 domain. The docking analyses predicted a more favorable interaction for L170R compared with wild-type GABPA, as reflected by a decreased weighted docking score. Notably, residue 170 was not located within the predicted GABPA-CBP/p300 binding interface, suggesting the observed differences are unlikely to arise from direct contacts with CBP/p300. Instead, the effect of the variant may be mediated through mutation-induced intramolecular rearrangements within GABPA that subtly alter its interaction surface. Detailed docking models together with the intermolecular interaction analyses are provided in **Supplemental Figure 3** and **Supplemental data file 3**.

### GABPA protein is detected in endothelial and mural cells of vascular tumor lesions

To visualize GABPA protein expression, tumor sections (left elbow, knee, and ankle) were stained with anti-human GABPA (magenta), the human endothelial marker Ulex europeus agglutinin-1 (UEA1) (yellow), and DAPI (cyan) for nuclei. Age-matched skin and proliferating phase IH were stained in parallel for comparison. Nuclear localized GABPA was detected in vessel-lining endothelial cells and in surrounding perivascular cells in all tumor lesions and in endothelial cells in IH and normal skin (**Figure 3G-I**). Quantification of GABPA-positive cells per total nuclei shows GABPA significantly increased compared to IH or to normal skin (**Figure 3J**). A limitation is that the anti-GABPA does not distinguish between GABPA-WT and GABPA-L170R.

### GABPA-L170R Alters Vascular Architecture in Zebrafish Embryos

We generated mosaic expression of wild-type and variant GABPA in zebrafish embryos using the endothelial-specific *fli1ep* promoter and the *cxcl12b* promoter, leading to expression in stromal cells (**Figure 4A**, **E**) and assessed vascular development under these conditions. The normal caudal aorta (CA), caudal vein plexus (CVP), and intersegmental vessels (ISV) can be easily discriminated at 48 hours post fertilization (hpf) using brightfield and fluorescence microscopy (**Figure 4B**, **F**). We assessed the malformation rate using empty vector controls, as an internal technical control, resulting in a malformation rate of 5-10% (**Supplemental Figure 4A, D, E, H**). Expression of wild-type GABPA under the control of either the *fli1ep* or *cxcl12b* promotor resulted in approximately 15-20% of embryos with a vascular malformation (**Supplemental Figure 4B, D, F, H**). Intriguingly, mosaic endothelial-specific expression of GABPA-L170R via the *fli1ep* promoter did not result in a significant increase of the malformation rate (**Figure 4C, D**). In contrast, mosaic stromal expression of GABPA-L170R via the *cxcl12b* promoter did cause a significant increase in the rate of vascular malformations to approximately 35-40% at 48 hpf (**Figure 4G, H**), resulting in both fusion of the caudal aorta and caudal vein plexus as well as aberrant ISVs. Together, these data implicate the GABPA-L170R variant as damaging to the developing vasculature and suggest that the variant may exert its disruptive effects via the stromal cell compartment.

**Figure 4:**
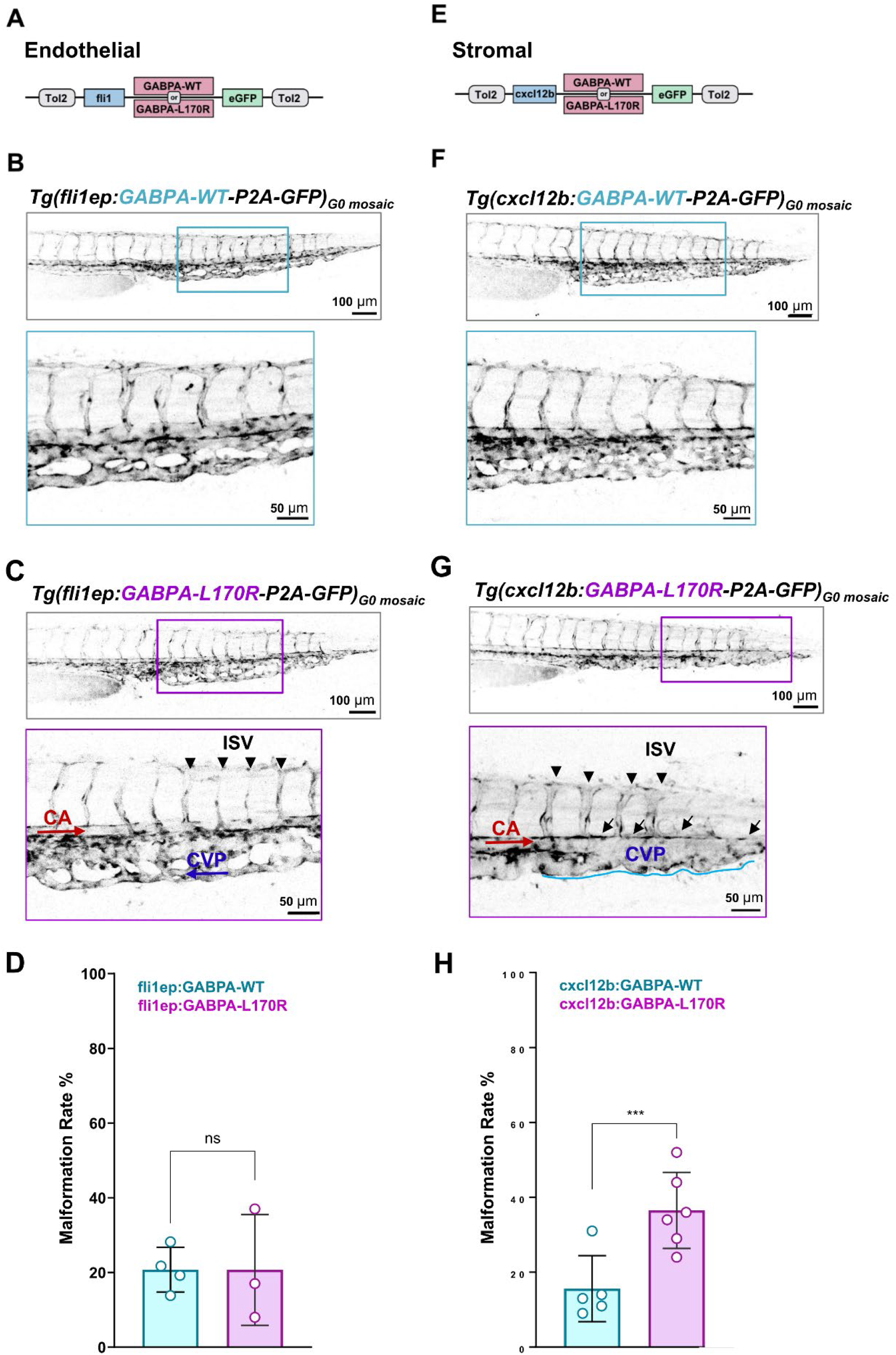
Expression of GABPA-L170R in a zebrafish model. (**A–D**) Functional evaluation of mosaic endothelial-specific wild-type versus mutant GABPA expression in zebrafish embryos. (**A**) Schematic of the transgenic construct driving either GABPA-WT or GABPA-L170R-expression linked via a self-cleaving P2A peptide to eGFP under control of the endothelial-specific *fli1ep* promoter. (**B, C**) Inverted fluorescence confocal images of the tail vasculature in transgenic embryos at 48 hours post-fertilization (hpf) expressing (**B**) *Tg(fli1ep:GABPA-WT-P2A-GFP)_G0 mosaic_* or (**C**) *Tg(fli1ep:GABPA-L170R-P2A-GFP)_G0 mosaic_*. Boxes framed in magenta and cyan indicate areas shown at higher magnification below. Intersegmental vessels (ISV, black arrowheads), caudal aorta (CA, red arrows), and caudal vein plexus (CVP, blue arrows) indicate globally intact and regular vascular patterning in both cohorts (scale bars: main panels, 100 µm; insets, 50 µm). (**D**) Quantification of the vascular malformation rate (%) following endothelial-specific expression shows no significant difference between groups. *n* = 3-4 independent experimental fish cohorts representing *n* = 133-191 individual zebrafish embryos; *ns* = not significant. (**E**-**H**) Functional evaluation of stromal wild-type versus mutant GABPA expression in zebrafish embryos. (**E**) Schematic of the transgenic construct driving gene expression under control of the stromal *cxcl12b* promoter. (**F**, **G**) Representative tail vasculature images of 48 hpf embryos expressing (**F**) *Tg(cxcl12b:GABPA-WT-P2A-GFP)_G0 mosaic_* or (**G**) *Tg(cxcl12b:GABPA-L170R-P2A-GFP)_G0 mosaic_*. Boxes framed in magenta and cyan indicate areas shown below at high magnification. Stromal expression of wild-type GABPA yields normal vessel architecture (**F**). In contrast, stromal expression of the p.L170R variant induced distinct developmental vascular anomalies marked by a fusion of the caudal aorta/caudal vein plexus (black arrows), a dilated venous plexus with non-directed flow, and enlarged, tortuous, or missing intersegmental vessels (ISV, black arrowheads) (**G**) (scale bars: main panels, 100 µm; insets, 50 µm). (**H**) Quantification shows significantly increased vascular malformation rate (%) in the group with stromal expression of the GAPBA variant compared to wild-type GABPA expressing controls. *p* = 0.0001; *n* = 5-6 independent fish cohorts; *n* = *299-341* individual fish. Fisher’s exact test. Data presented as mean ± SD. All depicted embryos are *Tg(kdrl:mCherry)*; the endothelial fluorescence signal is represented in inverted grayscale to clearly show the vascular architecture.

## SUMMARY

We identified a *de novo c.509T>G* variant in the alpha subunit of the transcriptional activator GABP in a child born with extensive vascular tumors covering her limbs and torso while sparing the face. The variant generates an amino acid substitution L170R in the PNT domain of GABPA, a protein-protein interaction domain characteristic of ETS family transcription factors. The *de novo c.509T>G* is absent from common reference human genome databases. Comparing the predicted intramolecular interactions in the PNT domains of GABPA-WT and GABPA-L170R shows that L170R substitution creates new electrostatic interactions in place of hydrophobic side chain interactions. Intermolecular interactions with its known transcriptional partner CBP/p300 appear stabilized by the GABPA-L170R variant. An amino acid substitution that alters protein-protein interactions between GABPA and its assisting transcriptional regulators could alter both the kinetics and the content of gene expression programs needed to achieve the appropriate levels of vascular cell differentiation, morphogenesis, and quiescence. Expression of the GABPA-L170R variant in an embryonic zebrafish model under the stromal promoter *cxcl12b* resulted in a significantly malformed developing vasculature, an effect that was not observed in the GABPA-WT-expressing embryo and, interestingly, also absent when GABPA-L170R was expressed under the endothelial-specific promoter *fli1ep*. This supports the pathogenic potential of GABPA-L170R with disruptive effects on vascular development arising in the stromal compartment.

Over the past decade, there has been immense progress made in understanding molecular genetics and biochemical pathways driving vascular anomalies (Goldenberg *et al*., 2025; Revencu *et al*., 2025; Seront *et al*., 2024). Transcriptional biology is an emerging focus in this field (Holm *et al*, 2025; Overman *et al*, 2019; Seebauer *et al*, 2022) and is rapidly advancing our understanding of the fundamental mechanisms of vascular development (Chen *et al*., 2025; Jeong *et al*., 2017). Herein, we identified a de novo variant in the ETS transcription factor GABPA that likely disrupts its interaction with associated key transcriptional co-activators. This discovery warrants further investigation into the role of GABPA in vascular development and vascular tumor pathology, particularly regarding how potential second somatic hits or local microenvironmental cues, like shear stress, drive this vascular tumor’s focal lesion presentation.

## METHODS

### Cell isolation and culture

Endothelial cells were isolated from vascular tumor tissue and expanded in culture on 1% gelatin-coated plates in Endothelial Basal Medium 131 (EBM131), 10% heat inactivated FBS, 2 mM glutamine, 100 U/ml penicillin, 100 mg/ml streptomycin, 2ng/ml bFGF as described (Boye *et al*., 2001).

### Histopathology and IF analysis

Histopathology was reviewed from 53 specimens obtained during 18 operative procedures between 3 months and 16 years of age in the Department of Pathology, Boston Children’s Hospital. As part of the routine diagnostic workup, immunohistochemical evaluation of several specimens was performed on formalin-fixed paraffin-embedded (FFPE) sections (5 μm) including H&E and immunostainings for CD31, CD34, PROX1, D2-40, and GLUT1 as part of the routine pathology workup (data not shown). Immunofluorescent stainings for CD31, UEA1, LYVE-1, GLUT1, and alpha-SMA following confocal imaging were performed on patient vascular tumor lesions from elbow, knee, and ankle as well as IH and normal skin tissues with three patients each (**Supplemental Figure 2A**). FFPE sections were deparaffinized and immersed in an antigen retrieval solution (citrate-EDTA buffer containing 10 mM citric acid, 2 mM EDTA, and 0.05% Tween-20, pH 6.2) for 20 minutes at 95°C-99°C. Sections were subsequently blocked for 30 minutes in 10% donkey serum (Abcam, ab7475) followed by incubation with a monoclonal anti-human CD31 antibody (1:30: Dako, 0823) or UEA1 directly labeled with Alexa Fluor 649 (1:50; Vector Laboratories, DL-1068-1) to stain for human endothelium. Primary antibody co-staining included polyclonal rabbit anti-human GABPA (1:100; Abcam, ab224325), monoclonal rabbit anti-human GLUT1 (1:100; Abcam, ab115730), monoclonal anti-human αSMA-Cy3 (1:500; Millipore Sigma, C6198, clone 1A4), and polyclonal rabbit anti-LYVE-1 (1:100; Abcam AB33682). Next, sections were incubated with secondary antibodies, including Alexa Fluor 488 donkey anti-rabbit IgG and Alexa Fluor 546 donkey anti-mouse or -rabbit IgG, respectively (1:200; Thermo Fisher Scientific, A21206, A10036, A10040). All slides were co-stained for DAPI to visualize nuclei and mounted thereafter (Thermo Fisher Scientific, R37606 and P36980). Immunofluorescent images were acquired by an LSM 980 confocal microscope (Zeiss) using a 20x or 63x objective lens. Images were analyzed using Fiji ImageJ software (version 1.54k, NIH, Bethesda, MD, USA). For perivascular wall thickness measurements in patient, IH, and normal skin control tissues, the distance from the outer edge of the alpha-SMA-positive cell layer to the UEA1-positive endothelium was manually measured (**Figure 2C–H**). GABPA was quantified manually by calculating the ratio of marker-positive nuclei to total DAPI-stained nuclei. For the patient, samples were derived from three distinct anatomical locations (left elbow, knee, and ankle); for IH and normal skin controls, specimens were obtained from n=3 independent individuals each. 5-10 representative images were captured at 20x magnification and quantified (**Figure 3J; Supplemental Figure 2A**).

### Genomic DNA isolation and Sequencing

Genomic DNA was isolated using the QIAamp DNA Micro kit (Qiagen, 56304). Genomic DNA quality control and fragment analysis were performed using the Agilent TapeStation platform at the Biopolymers Facility (Harvard Medical School, Boston, MA). DNAs extracted from blood of the patient and her parents were subjected to WES using the SureSelect v6 capture and Illumina paired-end reads of 2x100bp by Macrogen (Seoul, South Korea), now Psomagen. The average coverage without duplicates reached 246x for the index, 236x for the father and 243x for the mother. FFPE-DNA from lesions on the elbow and knee, as well as DNA of endothelial cells isolated from 7 locations (Boye *et al*., 2001) underwent WES using the SureSelect 8 capture with 2x150bp reads for an average coverage of 200x. In both cases, raw *FASTQ* files were aligned to the human reference genome hg38, followed by variant calling and import in the Highlander software (Brouillard *et al*, 2024). Sanger Sequencing on gDNA from HeomaEC-8b and HeomaEC-8g confirmed the presence of the *chr21_25752190 T>G* variant.

### Structural Modeling and Protein-Protein Docking

Structural analyses were performed using the AlphaFold-predicted structure of human GABPA (AF-Q06546-F1-model_v6). The L170R mutant model was generated by template-based homology modeling using the SWISS-MODEL server (https://swissmodel.expasy.org/), employing the wild-type GABPA structure as the template. Wild-type and mutant structures were subsequently analyzed using Discovery Studio Visualizer (v25. 1. 0. 24284) to assess mutation-induced changes in intramolecular interactions and local structural organization. To assess the potential impact of the L170R variant on CREBBP binding, protein-protein docking analyses were performed using both wild-type and L170R mutant GABPA structures against a truncated CREBBP fragment encompassing residues 1680-1850. This fragment was generated in PyMOL (v3.1.4.1; Schrödinger, LLC) from the AlphaFold-predicted CREBBP structure (AF-Q92793-2-F1-model_v6). Docking simulations were conducted using the ClusPro 2.0 protein-protein docking server (https://cluspro.org/). Docking models were ranked according to the ClusPro scoring algorithm, and he models with the lowest weighted scores, considering both cluster-center and lowest-energy solutions, were selected for further analysis. Predicted intra- and intermolecular interactions in the wild-type and mutant GABPA structures in interaction with CREBBP were examined using BIOVIA Discovery Studio Visualizer (v25. 1. 0. 24284).

### Zebrafish husbandry

Maintenance and breeding of zebrafish (Danio rerio) were performed in the Aquatic Core Facility for Zebrafish and Xenopus research (AquaCore) at Institute for Disease Modeling and Targeted Medicine under standard conditions. Only embryos up to 5 days post-fertilization were used. All experiments were performed in accordance with German laws for animal care and the Regierungspräsidium Freiburg.

### Plasmid preparation

Plasmids for zebrafish expression were designed using ApE-A plasmid editor (Davis & Jorgensen, 2022). Homo sapiens GABPA sequence was obtained from the online database Ensembl (Transcript ID: ENST00000400075.4). Minimal codon optimization for expression in *Danio rerio* was performed based on the IDT codon optimization tool, solely replacing rare codons with synonymous, higher-frequency alternatives before ordering pME:GABPA-p.L170R plasmid from Twist Bioscience (South San Francisco, CA, USA). p5E:fli1ep was a gift from Nathan Lawson (Addgene plasmid 31160); p5E:cxcl12b was cloned and tested by FGK in LZs laboratory (Boston Children’s Hospital and Harvard Medical School, Boston, USA); p3E:P2A-eGFP was a gift from Kristen Kwan & Christian Mosimann (Addgene plasmid 195967); pDest:Tol2A2 was a gift from Wolfgang Driever (University of Freiburg, Freiburg, Germany). Final expression constructs were cloned by Gateway Assembly using p5E:fli1ep or p5E:cxcl12b, pME:GABPA-p.L170R, p3E:P2A-eGFP and pDest:Tol2A2. The GABPA-WT variant of both constructs was cloned with Q5 Site-Directed Mutagenesis Kit (New England Biolabs, E0554S) with the following primer pair (Metabion): GABPA-WT-fwd (5′-GGCTGCTGCActgGAAGGCTATA-3′); GABPA-WT-rev (5′-CATCTTGTCACTTGTTCTGAAGTTTC-3′). Mutagenesis primers were designed using NEBaseChanger online tool. All plasmids were sequenced by Eurofins Genomics, whole plasmid sequencing service to confirm the final sequence.

### Transposase mRNA synthesis

pCS2FA-transposase plasmid, that contains the transposase gene under control of the SP6 promoter, was linearized with NotI-HF enzyme (New England Biolabs, R3189S) for 1 h at 37 °C. The digested sample was purified using the QIAquick PCR & Gel Cleanup Kit (Qiagen, 28506) according to the manufacturer’s protocol. Capped Tol2 transposase mRNA was synthesized from purified DNA using the mMESSAGE mMACHINE SP6 Transcription Kit (ThermoFisher Scientific, AM1340) according to the manufacturer’s protocol. The resulting mRNA was separated into 5 µl aliquots and stored at −20 °C to prevent freeze–thaw cycles.

### Zebrafish microinjection

A day before each injection, adult zebrafish pairs were transferred to small breeding tanks with a barrier separating male and female. Before the injection, the barrier was removed to allow spawning. The same day, 10 µl fresh injection mix was prepared with 20 ng/µl final concentration of the respective GABPA construct, 20 ng/µl final concentration of Transposase RNA, Phenol red solution (Sigma-Aldrich, 1072420) at 10% of the final volume, and nuclease-free water to a final volume of 10 µl. The injection mix was administered into *Tg(kdrl:mCherry)* embryos at the one-cell stage using Pneumatic PicoPump PV830 (World Precision Instruments) with an injection volume of 1 nl. For better readability, *Tg(kdrl:mCherry)* embryos injected with a construct containing the gene of interest (e.g. fli1ep:GABPA-WT-P2A-GFP or cxcl12b:GABPA-WT-P2A-GFP) are abbreviated as *Tg(fli:GABPA-WT-P2A-GFP)*_G0 mosaic_ or *Tg(cxcl12b:GABPA-WT-P2A-GFP)_G0 mosaic_*.

### Microscopy for zebrafish embryos

At 24 hpf, 1-phenyl 2-thiourea (PTU; 0,03 mg/ml working concentration) was added to Danieau’s media to prevent pigment formation and embryos were kept at 28 °C in an incubator. At 48 hpf, eGFP+ embryos were assessed for the presence of vascular malformations using a LEICA MZ10 F fluorescence microscope. Imaging plates for confocal microscopy were prepared in 35 mm glass bottom dishes with 1.5% agarose in Danieau’s media, 1-phenyl 2-thiourea (PTU; 0,03 mg/ml working concentration) and tricaine (0.168 mg/ml working concentration) mix using previously designed and 3D-printed molds (Kapp *et al*, 2024). Embryos were anesthetized with 0.168 mg/ml tricaine in Danieau’s media at room temperature and gently positioned laterally inside the trenches. Confocal microscopy images were acquired as z-stacks with ZEISS Celldiscoverer 7 with LSM 900. Images were obtained with the 488 nm and 561 nm lasers, 5X (NA 0.35) objective with a slice interval of 5.44 μm unless otherwise specified. Image processing performed with Fiji ImageJ software.

### Statistical analysis

Data were analyzed and plotted using GraphPad Prism 11.1.0 (Graph-Pad Software). Data are presented as mean ± SD for all experiments. Differences were considered significant for *P* values less than 0.05 with *P* *< 0.05, **< 0.01; ***< 0.001; **** < 0.0001. Figures were created using Affinity Designer 1.10.6. The malformation rate was calculated by dividing the number of eGFP-positive embryos exhibiting vascular malformations (e.g., fusion of the caudal aorta and the venous plexus) by the total number of eGFP-positive embryos. Statistical analysis was performed using two-tailed Fisher’s exact test of significance for malformation rate analysis.

### Study approval

Patient specimens and blood, as well as parents’ blood, were obtained under a protocol approved by the Committee on Clinical Investigation at Boston Children’s Hospital upon written informed consent of the patient’s guardian (IRB protocol number 04-12-175R).

## Acknowledgements

We extend our sincere gratitude to the patient and her family whose participation and support made this work possible. We thank the interdisciplinary team of the Vascular Anomalies Center at Boston Children’s Hospital for their support throughout this work.

Research reported in this manuscript was supported by a Pilot Study Award from Boston Children’s Hospital (2018) (to JB). AH was supported by the German Research Foundation (Walter Benjamin Fellowship, HO6799-1-1) and the Vascular Anomalies Center at Boston Children’s Hospital. AH, FGK, and MV are members of the Vascular Anomalies Working Group (VASCA) of the European Reference Network for Rare Multisystemic Vascular Diseases (VASCERN) - Project ID: 769036. Additionally, the studies were supported by the Fonds de la Recherche Scientifique - FNRS Grants T.0240.23 & P.C005.22 and by the Fund Generet managed by the King Baudouin Foundation (Grant 2018-J1810250-211305), the Walloon Region through the FNRS-WELBIO strategic research program (WELBIO IP X.1548.24), and the 21CVD03 Leducq Foundation Networks of Excellence Program grant “ReVAMP” (all to MV). PB is a Scientific Logistics Manager of the Genomics Platform of University of Louvain. The project was also supported by a Pierre M. fellowship. Imaging of the zebrafish embryos was performed at the ZEISS Celldiscoverer 7 at the Lighthouse Core Facility, funded in part by the Medical Faculty, University of Freiburg (Project Number: 2025/B3-Fol). We thank the staff of the aquatic core facility (AquaCore – RI00544) at the University Freiburg Medical Center – IMITATE, Germany, for the excellent support with the zebrafish maintenance and experimentation. FGK was supported by the charity cycling tour “Tour der Hoffnung” and received funding from the German Federal Ministry of Education and Research ((BMBF): EJPRD Joint Transnational Call 2023, NARRATIVE;FKZ 01GM2405) and the German Research Foundation (project number 571386042). We thank the Biopolymers Facility at Harvard Medical School (Boston, MA) for providing DNA quality control and fragment analysis services as well as the IDDRC Cellular Imaging Core (RRID: SCR026485), funded by NIH P50 HD105351, and S10 OD030322. The funders had no role in the study design, data collection and interpretation, or decision to submit this work for publication.

## Author contributions

AH: Conceptualization, Data Curation, Formal Analysis, Funding Acquisition, Investigation, Methodology, Project Administration, Visualization, Writing – Original Draft

PB: Formal Analysis, Investigation, Methodology, Software, Writing – Review & Editing

NM: Formal Analysis, Investigation, Methodology, Writing – Review & Editing

MM: Formal Analysis, Investigation, Methodology, Software, Writing – Review & Editing AS: Formal Analysis, Investigation, Methodology, Writing – Review & Editing

DB: Formal Analysis, Investigation, Methodology, Writing – Review & Editing

NL: Formal Analysis, Investigation, Methodology, Software, Writing – Review & Editing

AA: Investigation, Resources, Writing – Review & Editing

LIZ: Funding Acquisition, Supervision, Validation, Writing – Review & Editing

WE: Investigation, Resources, Writing – Review & Editing

HK: Investigation, Resources, Writing – Review & Editing

FGK: Funding Acquisition, Investigation, Methodology, Supervision, Validation, Writing – Review & Editing

MV: Funding Acquisition, Supervision, Validation, Writing – Review & Editing

JBM: Data Curation, Investigation, Resources, Writing – Review & Editing

JB: Conceptualization, Data Curation, Funding Acquisition, Supervision, Validation, Writing – Original Draft

