## Supplemental Figures 1-4 for "A de novo *GABPA* Variant in a Patient with Multifocal Cutaneous Vascular Tumors of an Unclassified Entity"

Holm A*, Brouillard P*, Mahammadzade N*, Mehrabipour M* et al.

| **Summary of Clinical Presentation** | **Multimodal Treatment Timeline** |
|---|---|
| Multiple congenital cutaneous vascular tumors:<br>• Ears and lower lips<br>• Extremities<br>• Trunk<br>• External genitalia, gluteal region<br><br>Functional impairment (growth):<br>• Left radial head dislocation<br>• Hip pain (bilateral)<br>• Hearing loss (left-sided)<br><br>Other findings:<br>• Pyogenic granuloma (left temple and cheek)<br>• Arachnoid cyst (posterior fossa) | MEDICAL THERAPY:<br><br>5 – 7/1997: **Systemic corticosteroids**<br>No response: Growth and occurrence of new lesions*<br><br>7/1997 – 4/1998: **Interferon alpha-2a**<br>Partial stabilization; discontinued (spastic diplegia)<br><br>6 – 10/1999: **Thalidomide**<br>No response*<br><br>11/1999 – 3/2000: **Cyclophosphamide** and **Vincristine**<br>No response*<br><br>SURGERY:<br><br>9/1997 – 11/2013: **Multiple surgical excisions** (27):<br>Partial regrowth of large lesions<br><br>3/1999: Left posterior fossa craniotomy:<br>Partial improvement of spastic diplegia<br><br>9/1998: Excision of pyogenic granulomas<br><br>LONG-TERM OUTCOMES:<br><br>As of 2005: Stabilization, softening, and spontaneous regression<br><br>As of 2016: New punctuate, flat, bluish cutaneous lesions (upper extremities) |

**Supplemental Figure 1.** Summary of clinical presentation and treatment course. Summary of patient records during her treatment by Dr. John B. Mulliken and his team at the Vascular Anomalies Center at Boston Children's Hospital.

**A**

| Diagnosis | Sex | Age at Resection | Anatomical Location/Source |
|---|---|---|---|
| **Diffuse Vascular Tumors** | Female | 8 months – 8 years | Left elbow, knee, and ankle |
| **Infantile Hemangioma** | Female | 3 months | Lower occipital scalp |
| | Female | 5 months | Left posterior scalp |
| | Female | 7 months | Right abdominal wall |
| **Normal Skin Control** | Male | 8 months | Foreskin |
| | Male | 1 year | Foreskin |
| | Female | 7 years | Forearm margin (Nevus excision) |

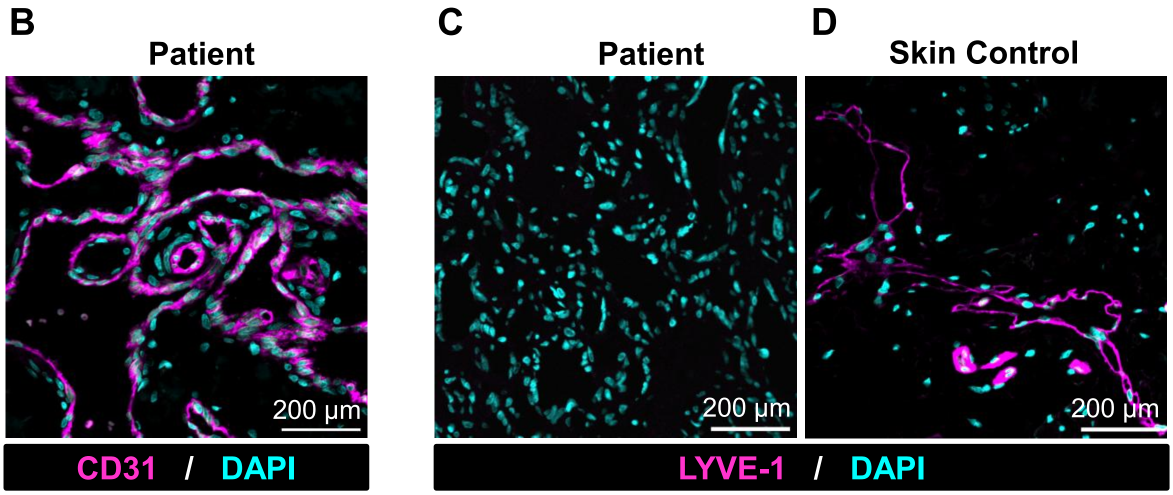


**Supplemental Figure 2.** Endothelial marker expression and patient characteristics of tissue specimen. (**A**) Immunofluorescence image of patient tumor tissue (left knee) stained with anti-CD31, a pan endothelial marker (magenta). (**B**) Patient's tumor section stained for with anti-human LYVE-1, a lymphatic endothelial marker (magenta). (**C**) Normal age-matched skin stained in parallel as a positive control in. Cell nuclei are co-stained with DAPI (cyan). Scale bars: 200 µm. (**D**) Summary of patient characteristics of tissue samples used throughout the study.

**A**

| Sample | Location | Coverage (reads) | Reference (T) | Alternative (G) | VAF |
|---|---|---|---|---|---|
| *Blood* | | | | | |
| Mother's blood | | 187 | 187 | 0 | 0.00% |
| Father's blood | | 178 | 178 | 0 | 0.00% |
| Patient's blood 1 | | 199 | 112 | 87 | 43.72% |
| Patient's blood 2 | | 110 | 50 | 60 | 54.55% |
| *Patient Endothelial Cells* | | | | | |
| HeomaEC-8a | L elbow | 142 | 73 | 69 | 48.59% |
| HeomaEC-8b | L ankle | 119 | 60 | 59 | 49.58% |
| HeomaEC-8c | L knee | 130 | 79 | 51 | 39.23% |
| HeomaEC-8f | R medial foot | 142 | 76 | 66 | 46.48% |
| HeomaEC-8g | R thigh | 106 | 56 | 50 | 47.17% |
| HeomaEC-8h | R dorsal foot | 124 | 68 | 56 | 45.16% |
| HeomaEC-8i | R distal tibia | 121 | 74 | 47 | 38.84% |
| *Patient tumor tissue (FFPE)* | | | | | |
| L elbow | | 96 | 63 | 33 | 34.38% |
| L knee | | 56 | 25 | 31 | 55.36% |

**B**

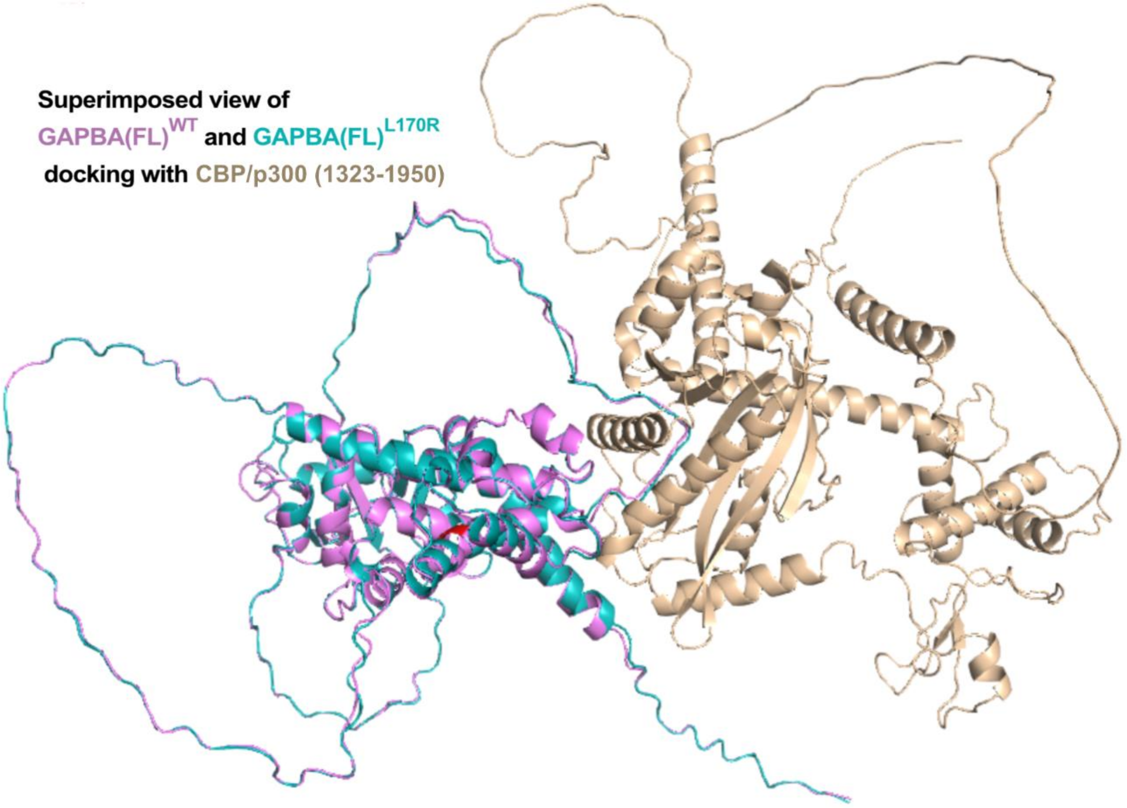


| | Representative | Weighted Score |
|---|---|---|
| GAPBA(FL)$^{WT}$ vs CBP/p300 (1323-1950) | Center | -1274.3 |
| | Lowest Energy | -1358.4 |
| GAPBA(FL)$^{L170R}$ vs CBP/p300 (1323-1950) | Center | -1310.9 |
| | Lowest Energy | -1367.8 |

Supplemental Figure 3

**Supplemental Figure 3.** Legend of cells and tissue used for WES and docking analysis of GABPA-WT and GABPA-L170R with CBP/p300. (A) Summary of blood and tissue samples from the patient and her parents underlying the WES, including location, coverage, reference, alternative, and variant allelic frequencies (VAF). (B) Structural superimposition of docking complexes generated using full-length GABPA and a CBP/p300 fragment encompassing residues 1323-1950. Wild-type GABPA and L170R-GABPA are shown in magenta and cyan, respectively, whereas CBP/p300 is shown in wheat. The L170R variant exhibited a more favorable docking score compared with wild-type GABPA. Residue 170 (red) was not located within the predicted GABPA-CBP/p300 binding interface in either complex, supporting an indirect effect of the variant on CBP/p300 binding. Detailed intermolecular and intramolecular interaction maps are provided in **Supplemental figure 3, data file**.

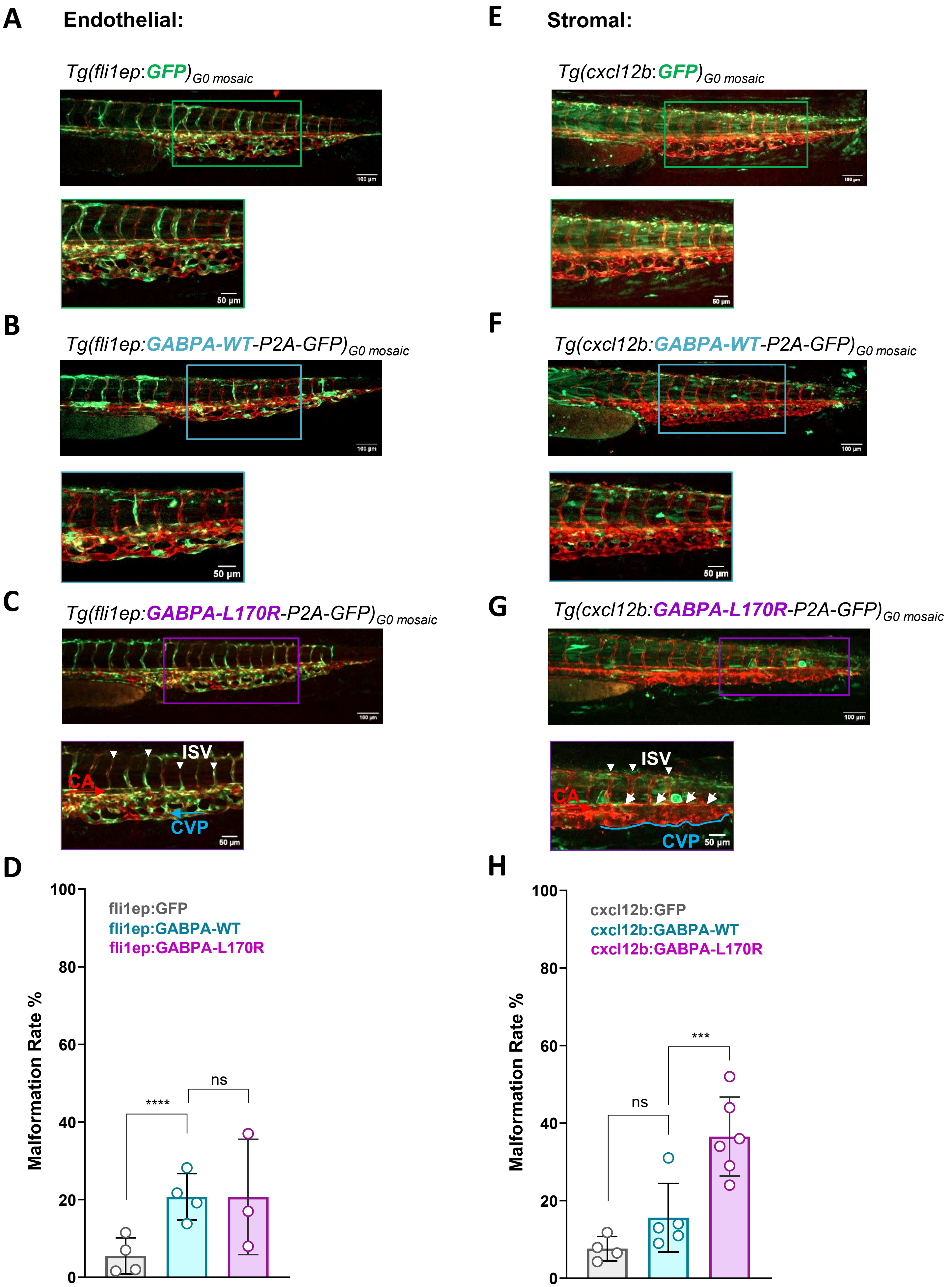


Supplemental Figure 4

**Supplemental Figure 4.** Imaging of vasculature in zebrafish model. Zebrafish vascular phenotype with assigned pseudo-colors. Red color assigned for mCherry channel (depicting Tg(kdrl:mCherry)) and green color for eGFP channel (depicting the mosaically-expressed constructs). Boxed areas are shown at higher magnification below. (A) Tg(fli1ep: GFP)G0 mosaic (empty vector control), Tg(fli1ep:GABPA-WT-P2A-GFP)G0 mosaic, and Tg(fli1ep:GABPA-L170R-P2A-GFP)G0 mosaic show normal vascular architecture of the tail vasculature at 48 hpf. (C) Tg(cxcl12b: GFP)G0 mosaic (empty vector control) and Tg(cxcl12b:GABPA-WT-P2A-GFP)G0 mosaic show normal vascular architecture of the tail vasculature at 48 hpf, while the vasculature in Tg(cxcl12b:GABPA-L170R-P2A-GFP)G0 mosaic shows the abnormalities described in Figure 4 (fusion of artery and vein, dilation of the venous plexus, distorted intersegmental vessels (ISV)). Labels in the lowest panel show selected ISV (white arrowheads), the caudal aorta (CA, red arrow), and the caudal vein plexus (CVP, blue arrow), with the arrows also indicating the direction of blood flow. Scale bars 100 μm and 50 μm. (B and D) Quantification of malformation rate in zebrafish embryos. (B) Quantification for embryos injected with fli1ep:GFP empty vector control with malformation rates found in fli1ep:GABPA-WT and fli1ep:GABPA-L170R injected embryos, the last two as depicted in Figure 4. $p < 0.0001$ for fli1ep:GFP vs fli1ep:GABPA-WT; ns = not significant. n = 3-4 independent fish cohorts; n = 133-279 individual fish. Fisher's exact test. Data presented as mean ± SD. (D) Quantification for embryos injected with cxcl12b:GFP empty vector control with malformation rates found in cxcl12b:GABPA-WT and cxcl12b:GABPA-L170R injected embryos, the last two as depicted in Figure 4. ns = not significant. n = 4-6 independent fish cohorts; n = 292-341 individual fish. Fisher's exact test. Data presented as mean ± SD.
